# SARS-CoV-2 variant B.1.1.7 caused HLA-A2^+^ CD8^+^ T cell epitope mutations for impaired cellular immune response

**DOI:** 10.1101/2021.03.28.437363

**Authors:** Chanchan Xiao, Lipeng Mao, Zhigang Wang, Guodong Zhu, Lijuan Gao, Jun Su, Xiongfei Chen, Jun Yuan, Yutian Hu, Zhinan Yin, Jun Xie, Weiqing Ji, Haitao Niu, Feng Gao, Oscar Junhong Luo, Lianbo Xiao, Pengcheng Wang, Guobing Chen

## Abstract

The rapid spreading of the newly emerged SARS-CoV-2 variant, B.1.1.7, highlighted the requirements to better understand adaptive immune responses to this virus. Since CD8^+^ T cell responses play an important role in disease resolution and modulation in COVID-19 patients, it is essential to address whether these newly emerged mutations would result in altered immune responses. Here we evaluated the immune properties of the HLA-A2 restricted CD8^+^ T cell epitopes containing mutations from B.1.1.7, and furthermore performed a comprehensive analysis of the SARS-CoV-2 specific CD8^+^ T cell responses from COVID-19 convalescent patients and SARS-CoV-2 vaccinees recognizing the ancestral Wuhan strain compared to B.1.1.7. First, most of the predicted CD8^+^ T cell epitopes showed proper binding with HLA-A2, while epitopes from B.1.1.7 had lower binding capability than those from the ancestral strain. In addition, these peptides could effectively induced the activation and cytotoxicity of CD8^+^ T cells. Our results further showed that at least two site mutations in B.1.1.7 resulted in a decrease in CD8^+^ T cell activation and a possible immune evasion, namely A1708D mutation in ORF1ab_1707-1716_ and I2230T mutation in ORF1ab_2230-2238_. Our current analysis provides information that contributes to the understanding of SARS-CoV-2-specific CD8^+^ T cell responses elicited by infection of mutated strains or vaccination.

**Graphical Abstract:** 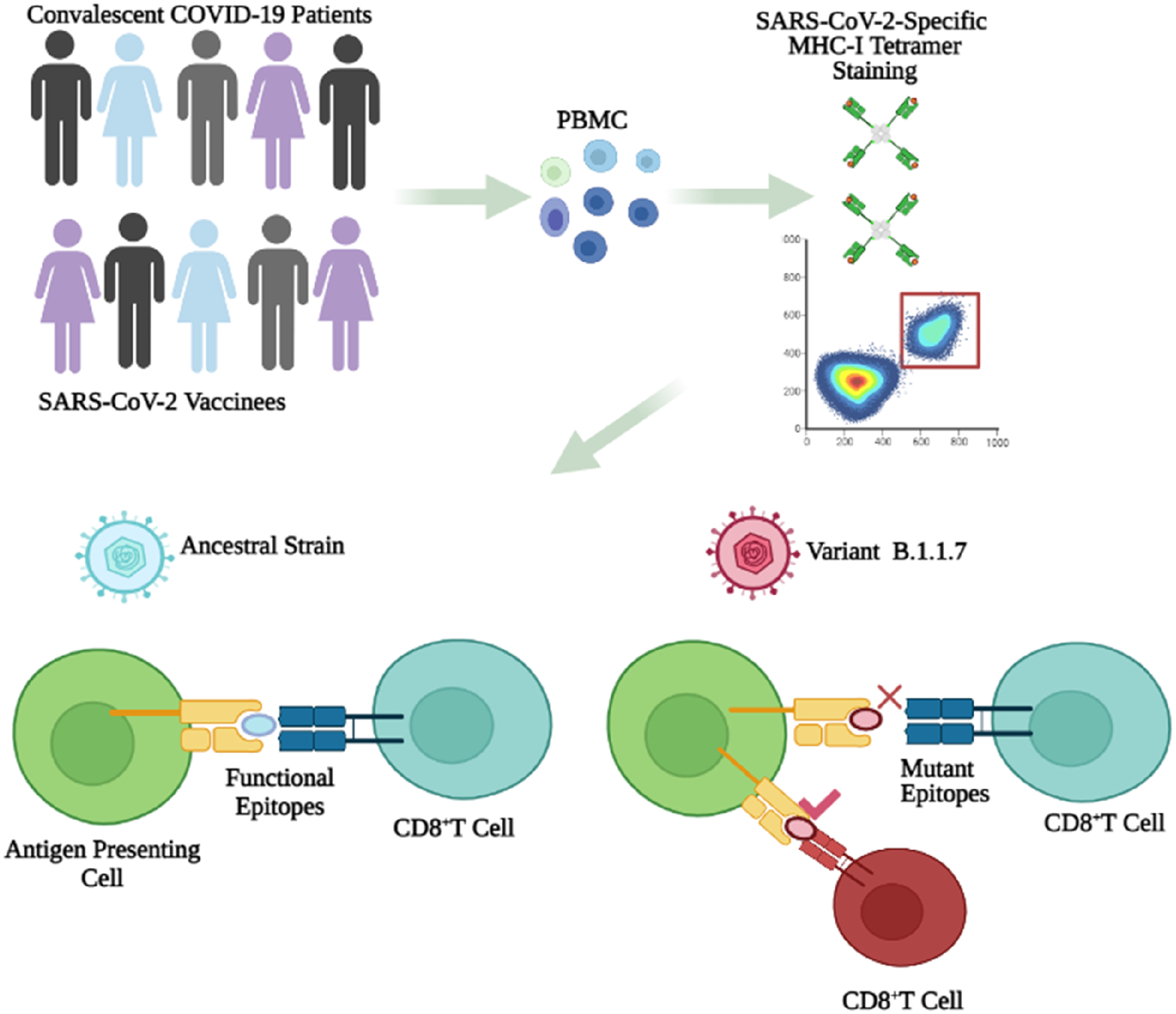

## INTRODUCTION

The coronavirus disease 2019 (COVID-19) pandemic has been sweeping the world. Its etiological agent, severe acute respiratory syndrome coronavirus 2 (SARS-CoV-2) belongs to the *Betacoronovirus* genus of the *Coronoviridae* family, and this single-stranded positive-sensed RNA virus bears 11 protein-coding genes, including 4 for structural proteins: spike (S), envelope (E), membrane (M) and nucleocapsid (N), and 7 for nonstructural proteins: open reading frame (ORF) 1ab, ORF 3a, ORF 6, ORF 7a, ORF 7b, ORF 8 and ORF 10(Wu et al., 2020). It’s believed that the viral clearance in SARS-CoV-2 infected individuals is mainly dependent on host immune system, especially adaptive immunity(Zhang et al., 2020). Specific antibodies have been observed in virus infected individuals and convalescent COVID-19 patients, with S and N being the major viral proteins to elicit antibody production(Wheatley et al., 2021). S protein bears the binding site to angiotensin converting enzyme 2 (ACE2) receptors on host cells and is crucial for viral infection. So the neutralizing antibodies against S protein are believed to play an important role for the virus clearance(Bertoglio et al., 2021). However, a couple of studies have shown that antibody titres decline fast in some convalescent patients, indicating a short duration of humoral immunity(Sette and Crotty, 2021). On the other hand, current studies have demonstrated that specific T cell responses emerge in most of the COVID-19 patients during the early stage of the infection(Ferreras et al., 2021). Although significant reduction in T cell counts was observed in severe COVID-19 patients, the revealed antigen specific T cell response indicated their important role in resolving SARS-CoV-2 infection(Grifoni et al., 2020; Le Bert et al., 2020) (Weiskopf et al., 2020). Furthermore, SARS-CoV-2 specific CD8^+^ T cells have been detected in convalescent COVID-19 patients(Braun et al., 2020; Grifoni et al., 2020) (Sattler et al., 2020) and SARS-CoV-2 vaccinees(Jackson et al., 2020). Recent studies have shown that specific CD8^+^ epitopes to SARS-CoV-2 are mainly located in ORF1ab, N protein, S protein, ORF 3, M protein and ORF 8(Ferretti et al., 2020; Grifoni et al., 2020), and the identification of these epitopes will provide the basis for next-generation vaccine development and better understanding of CD8^+^ T cell immunity.

With the ongoing spreading of the virus all over the world, the genetic evolution in SARS-CoV-2 continues to provide the opportunities for the virus to obtain mutations which might contribute to the changes in viral transmissibility, infectivity, pathogenesis and even immune evasion(Neches et al., 2021; Rashid et al., 2021). The D614G spike variant emerged in March 2020 was the earliest evidence of adaptive evolution of the virus in humans, which resulted in increased infectivity of the virus(Yurkovetskiy et al.). Recently, a newer variant termed B.1.1.7 (also called VUI202012/01) was spreading rapidly in the United Kingdom (UK) and raised great concerns(Davies et al., 2021; Kirby, 2021). This variant contains 17 non-synonymous mutations in ORF1ab, S protein, ORF8 and N proteins, some of which are of particular concerns, such as the D614G mutation and 8 additional mutations in S protein: ΔH69-V70, ΔY144, N501Y, A570D, P681H, T716I, S982A, and D1118H (Davies et al., 2021). For example, N501Y is located in the receptor binding motif (RBM) and P681H is proximal to the furin cleavage site(Peacock et al., 2020; Starr et al., 2020). ΔH69/ΔV70 deletion in S protein has evolved in other lineages of SARS-CoV-2, which enhances viral infectivity *in vitro* and is linked to immune escape in immunocompromised patients(Kemp et al., 2020a; Kemp et al., 2020b). There is strong evidence that variant B.1.1.7 is spreading substantially faster than preexisting SARS-CoV-2 variants(Davies et al., 2021; Kemp et al., 2020b; Volz et al., 2021). The model analysis suggests that this difference could be explained by an overall higher infectiousness of variant B.1.1.7. However, it is not clear that this is due to the shorter generation time or immune escape(Davies et al., 2021). Mutations in immune dominant epitopes might potentially alter their immunogenicity, and subsequently the immune responses of the host.

Our previous work has shown that the mutations in given CD8^+^ T cell epitopes resulted in antigen presentation deficiency and impaired antigen specific T cell function, indicating an immune evasion induced by viral evolution(Qiu et al., 2020). In this work, we predicted the potential CD8^+^ T cell epitopes within the areas where these mutations are located, and compared the immune properties of the ancestral and mutant peptides, including MHC I binding and activation of CD8^+^ T cells. Furthermore, we detected the epitope specific CD8^+^ T cells in convalescent COVID-19 patients and SARS-CoV-2 vaccinees by using corresponding tetramers. The results showed that at least two site mutations in the variant B.1.1.7 resulted in a decrease in CD8^+^ T cell activation and a possible immune evasion, namely A1708D mutation in ORF1ab_1707-1716_ and I2230T mutation in ORF1ab_2230-2238_. Our current analysis provides useful information that helps for better understanding of the SARS-CoV-2-specific CD8^+^ T cell responses elicited by infection of mutated strains or vaccination.

## RESULTS

### Identification of potential T cell epitopes containing mutations in B.1.1.7

During late 2020, the World Health Organization (WHO) announced the emergence of a novel coronavirus variant B.1.1.7 in the UK. We immediately carried out HLA-A2-restricted T cell epitope screening and identification of all the possible peptides containing the mutations in B.1.1.7 by using the high-throughput screening platform and artificial antigen presentation system for epitopes (Figure 1A-B, Supplementary Table 1). To validate these predicted epitopes, we first checked whether they could be presented by HLA-A2 on the antigen presenting cells (APC). T2A2 is an APC with TAP deficiency and HLA-A2 expression on cell surface. The peptide-MHC complex would be more stabilized if the epitopes bind with HLA-A2 suitably. Compared with the negative control peptide GLQ (GLQRLGYVL) from Zika virus, most of the predicted SARS-CoV-2 epitopes showed reasonable HLA-A2 binding. However, the HLA-A2 binding of most of the epitopes from the variant B.1.1.7 was reduced (Figure 1C-D). We further checked the direct binding of these epitopes to the purified HLA-A2 protein. According to the UV exchanged peptide-MHC assay, all the peptides, except for N S235F, exhibited strong binding to HLA-A2 (Figure 1E). The results indicated that majority of the predicted epitopes could form peptide-MHC complex (pMHC), and corresponding tetramers could be constructed next.

**Figure 1:**
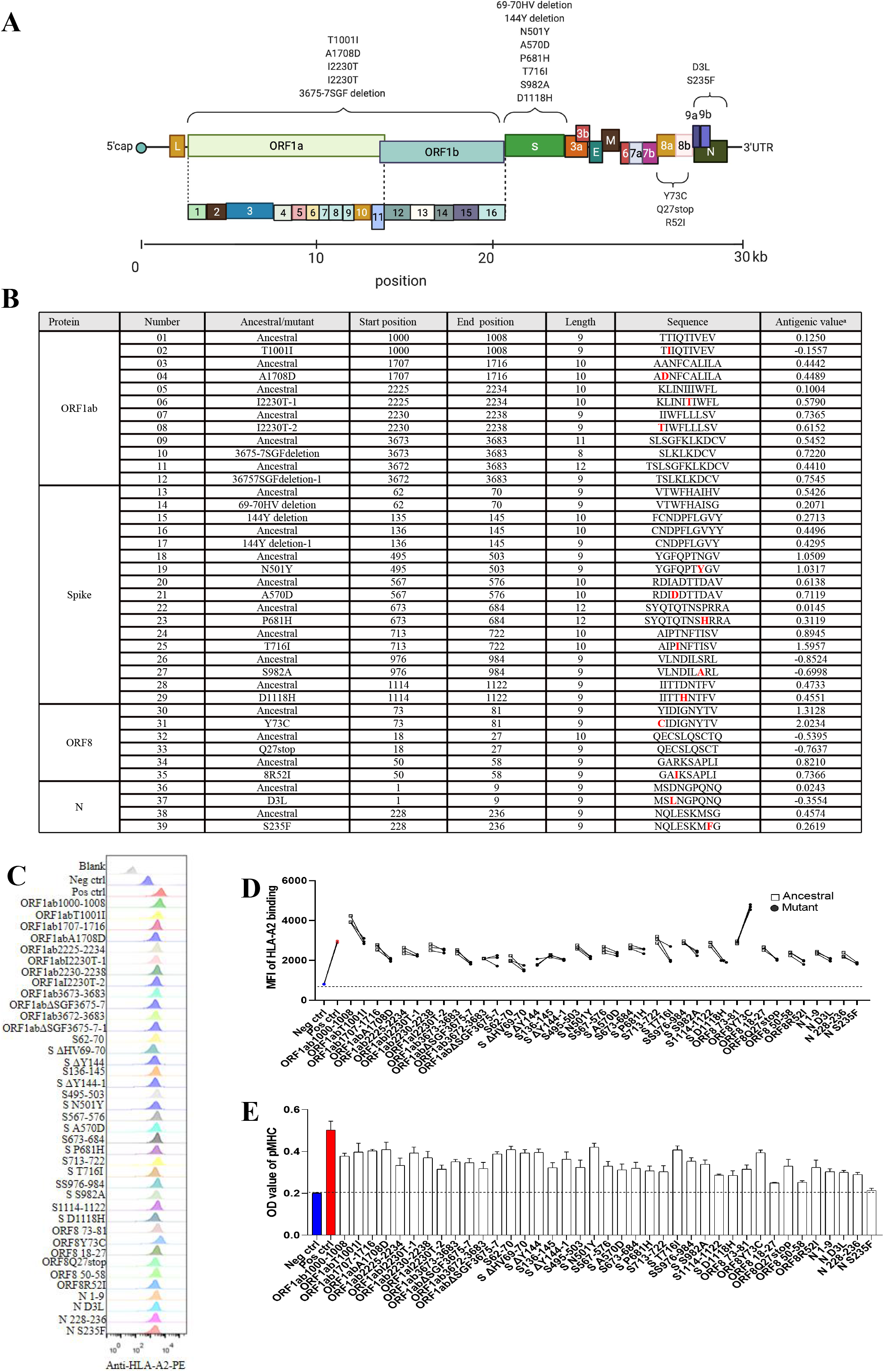
Identification of HLA-A2 restricted T cell epitopes on SARS-CoV-2 variant B.1.1.7. **A:** The schematic of mutation sites of the SARS-CoV-2 variant B.1.1.7. **B:** List of the predicted epitopes for following experiments. The mutated amino acids were highlighted as red in varian B.1.1.7t. ^a^The antigenic vaule threshold was > 0.4000 **C**-**D:** Comparison of ancestral and mutant epitope binding affinity to HLA-A2 in T2A2 cells. ancestral and mutated epitopes were synthesized and incubated with T2A2 cells. The binding of the peptide on T2A2 was measured with anti-HLA-A2 staining with flow cytometry. Binding capacity was presented as mean fluorescence intensity (MFI) of HLA-A2 staining. C was the representative plot of D. Each symbol represents an independent experiment. Ancestral: Wuhan strain epitope; Mutant: varian B.1.1.7t epitope. **E:** Evaluation of epitope binding to HLA-A2 with ELISA assay. Peptide exchanged assay was performed with coated UV-sensitive peptide/MHC complex and given peptides. The binding capability was measured with pMHC ELISA assay. Data shown are mean plus standard error of the mean (SEM). Threshold for pMHC formation positivity was set as above the average OD value of the negative-control cohort. Blank: no peptides; Neg ctrl: negative control, Zika virus peptide GLQRLGYVL; Pos ctrl: positive control, influenza A M1 peptide GILGFVFTL.

### Activation and cytotoxicity of T cells stimulated with T cell epitopes containing mutations of B.1.1.7

To further analyze whether the epitope-bound T2A2 cells could activate T cells, we tested the expression level of T cell activation marker CD69 and the proportion of peptide-specific CD8^+^ T cells after stimulation with peptide-bound T2A2 cells. As shown in Figure 2A-B, T2A2 cells bearing peptides of ancestral and B.1.1.7 induced significant increase in CD69 expression, respectively (Figure 2A-B). Further, our results showed that two mutations, namely A1708D in peptide ORF1ab_1707-1716_ and I2230T in peptide ORF1ab_2230-2238_ induced dramatically less proportion of specific CD8^+^ T cells than ancestral in the same subject. Intriguingly, the I2230T mutation in peptide ORF1ab _2225-2234_ did not result in similar decrease in T cell activation (Figure 2C-E) (Supplementary Figure 1 A-C). Next, we performed cross-detection of ancestral peptide specific CD8^+^ T cells with tetramers containing mutant peptides, and vice versa. The results showed that CD8^+^ T cells stimulated with the mutant peptides could not be recognized by tetramers containing the ancestral peptides (Figure 2F-G), nor could CD8^+^ T cells stimulated with the ancestral peptides be recognized by tetramers containing the mutant peptides (Figure 2H-I). Furthermore, both ancestral and mutant peptide-bound T2A2 cells stimulated T cell-mediated T2A2 killing. However, the mutant peptide group had a higher proportion of survival target cells compared to the ancestral group, suggesting a decreased cytotoxicity from mutant peptide specific CD8^+^ T cells (Figure 2J-K). In addition, the proportion of CFSE-Annexin V^+^ T2A2 was less for mutant peptide group than that for ancestral peptide group, indicating less T cell-mediated target cell apoptosis (Figure 2L-M). Finally, the CD8^+^ IFN-γ (Figure 2N-O) and Granzyme B (Figure 2P-Q) levels in mutant peptide group were significantly lower than those in ancestral peptide group. All the above results suggested that, compared to the ancestral, T cell mediated immune responses induced by B.1.1.7 mutant peptides were impaired.

**Figure 2:**
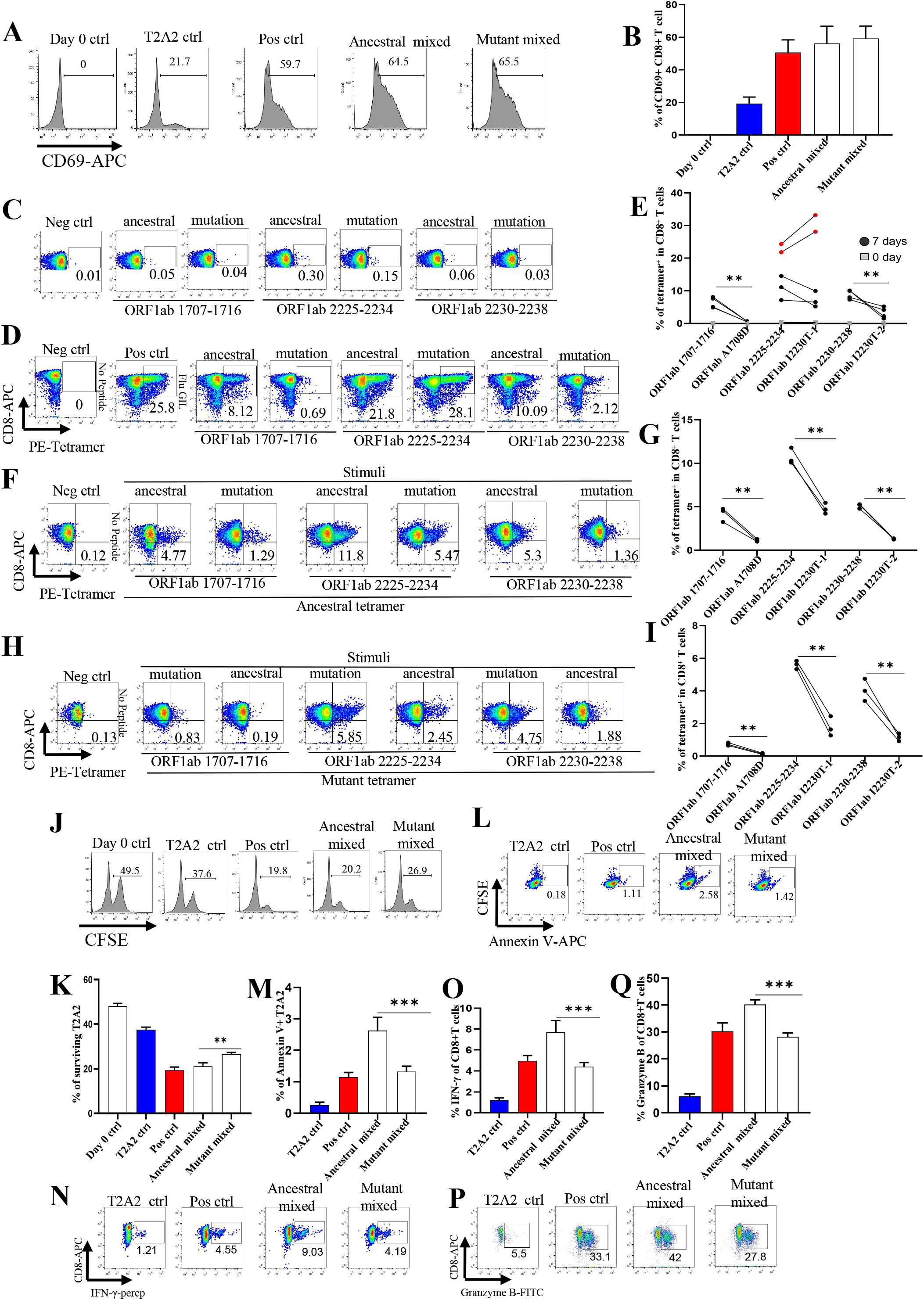
Evaluation of impaired immune protection caused by epitope mutation in SARS-CoV-2 variant B.1.1.7. Mitomycin pretreated T2A2 cells were loaded with mixed peptides from ancestral or mutant, and incubated with CD8^+^ T cell from health donors at 1:1 ratio, respectively. Epitope specific CD8^+^ T cells were generated after 7 day stimulation. **A**-**B:** The CD69 expression level of activated CD8^+^ T cell was evaluated with flow cytometry after 16 hour stimulation. A was the representative plot of B. n=4 per group. Day 0 ctrl: staining before stimulation; T2A2 ctrl: T2A2 without peptide loading; Pos ctrl: T2A2 loaded with influenza A M1 peptide GILGFVFTL. **C**-**E:** Epitope specific CD8^+^ T cell measurement before (C) and after (D) 7 day stimulation. The cells were stained with corresponding ancestral or mutated tetramer, and compared before and after stimulation (E). Four and five repeats were performed for decreased and nonchanged comparison, respectively. Please also see Supplementary Figure 1 A-C. **F-I:** Cross-detection of epitope specific CD8^+^ T cells with tetramers based on ancestral and corresponding mutant peptides. F-G: ancestral or mutant epitopes stimulated CD8^+^ T cells were stained with ancestral peptide-based tetramer. H-I: mutant or ancestral epitopes stimulated CD8^+^ T cells were stained with mutant peptide-based tetramer. n=3 per group. Symbols in G and I represented individual person. The *p* values were calculated by paired-samples T test. \*\**p* < 0.01. Neg ctrl: T2A2 without peptide loading; Pos ctrl: T2A2 loaded with influenza A M1 peptide GILGFVFTL. **J**-**M:** Epitope specific CD8^+^ T cell mediated cytotoxicity was evaluated after 7 day culture (J&K). The remained CFSE labeled T2A2 cells were calculated as survived target cells. J was the representative plot of K. Apoptosis of T2A2 cells at day 7 after culture. The proportion of CFSE^+^ AnnexinV^+^ cells was calculated as indicator for epitope stimulated T cell mediated T2A2 apoptosis (L&M). L was the representative plot of M. n=4 per group. **N**-**O:** The expression of IFN-γ after epitope stimulation for 7 days. IFN-γ was measured with intracellular stained flow cytometry. N was the representative plot of O. n=4 per group. **P**-**Q:** The expression of Granzyme B after epitope stimulation for 7 days. Granzyme B was measured with intracellular stained flow cytometry. P was the representative plot of Q. n=4 per group. Day 0 ctrl: staining before stimulation; T2A2 ctrl: T2A2 without peptide loading; Pos ctrl: T2A2 loaded with influenza A M1 peptide GILGFVFTL. The *p* values were calculated by paired-samples T test, \*\*\**p* < 0.001.

### SARS-CoV-2-specific CD8^+^ T cell profiling in convalescent COVID-19 patients and SARS-CoV-2 Vaccinees

We recruited a cohort of 25 convalescent COVID-19 patients and 60 SARS-CoV-2 vaccinees, including 4 and 17 who were HLA-A2 positive, respectively. The demographic and clinical information of the patients with tetramer^+^ cells were presented in Supplementary Table 2. We examined the *ex vivo* phenotypes of SARS-CoV-2 tetramer^+^ CD8^+^ T cells in PBMC of the patients by assessing the expression levels of the chemokine receptor CCR7 and CD45RA. It’s observed that the tetramers prepared with the above identified epitopes could recognize the specific memory T cells in convalescent patients (Figure 3A-B). MHC class I tetramer+ cells predominantly exhibited an effector memory (CCR7^-^CD45RA^-^) phenotype, and roughly 20% of tetramer^+^ cells were naïve T (CCR7^+^CD45RA^+^) cells (Figure 3C-D). Meanwhile, in the same individual, the proportion of T cells recognized by the B.1.1.7 mutant epitope tetramers was lower than that by the ancesral epitope tetramers (Figure 3B). We then examined the differentiation phenotype of SARS-CoV-2 specific CD8^+^ T cells in vaccinees, with the information of the subjects presented in Supplementary Table 3. It’s observed that the tetramers prepared with the above identified epitopes could recognize the specific memory T cells (Figure 3E-G), and the majority of tetramer^+^ cells were terminally differentiated effector (CCR7^-^CD45RA^+^) cells (Figure 3H-I). Similarly, the proportion of T cells recognized by the B.1.1.7 mutant epitope tetramers was lower than that by the ancestral epitope tetramers in the same individual (Figure 3F). All above data indicated that these emerged mutations might have caused a deficiency in the antigen presentation of the dominant epitopes, which was required to rebuild a new CD8^+^ T cell immune response in COVID-19 patients and vaccinees.

**Figure 3:**
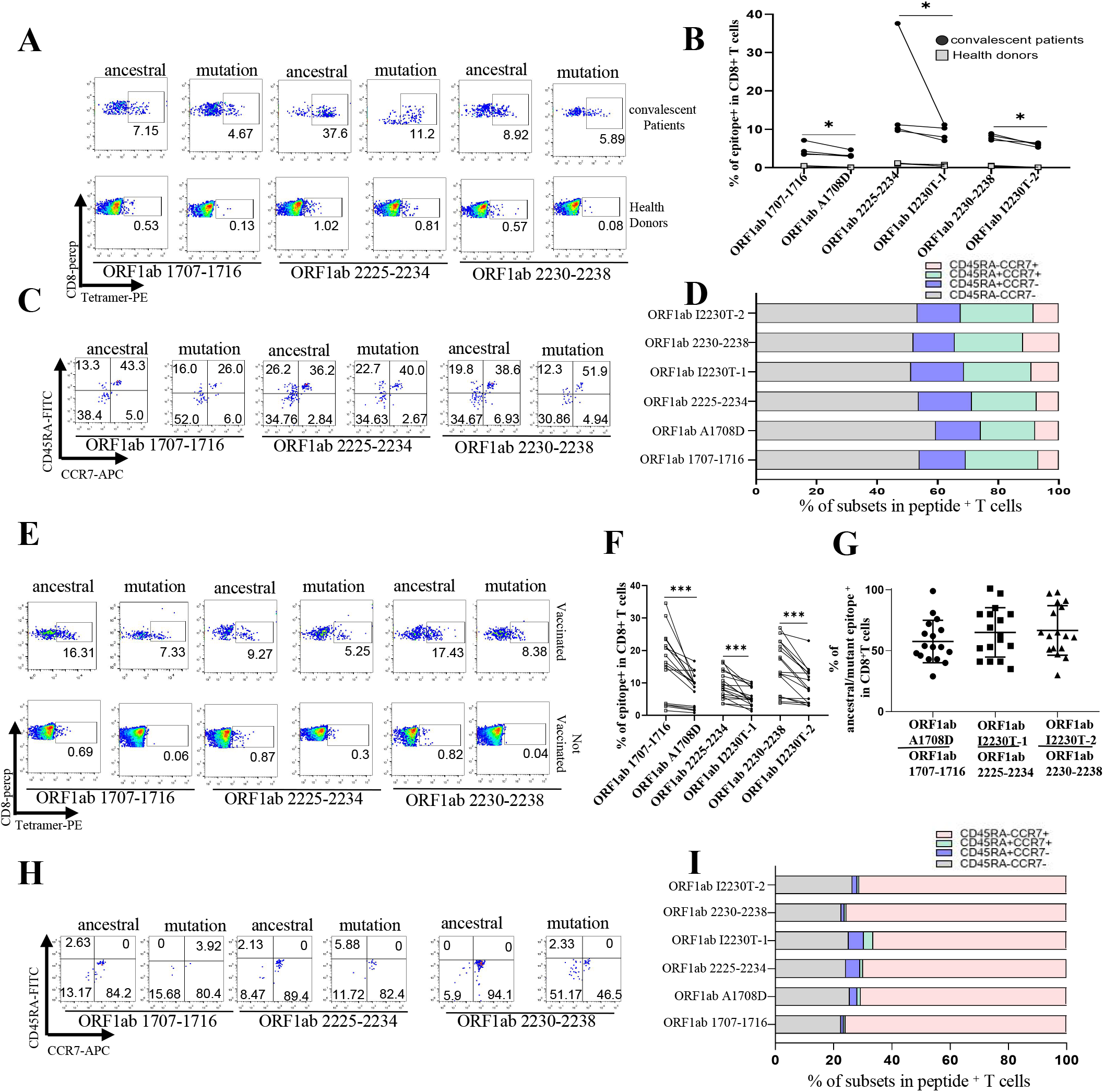
Profiling of epitope specific CD8^+^ T cells in convalescent COVID-19 patients and SARS-CoV-2 vaccinees. **A-D:** Ancestral and mutant peptide specific CD8^+^ T cells (A-B) and functional subtypes (C-D) in HLA-A2^+^ convalescent COVID-19 patients and healthy donors were measured with flow cytometry. Ancestral and mutant epitope specific CD8^+^ T cells in the same individual were compared in B. n=4 per group. **E-I:** Peripheral venous blood was collected from HLA-A2^+^ vaccinees 1-3 months after SARS-CoV-2 vaccination. Ancestral and mutant peptide specific CD8^+^ T cells (E-G) and functional subtypes (H-I) were measured with flow cytometry. The ancestral and mutant epitope specific CD8^+^ T cells in the same individual were compared in F, with ration calculation in G. n=17 per group. The *p* values were calculated by paired-samples T test. \**p* < 0.05, \*\**p* < 0.01, \*\*\**p* < 0.001.

### Computational molecular docking simulation of ancestral and B.1.1.7 epitopes with HLA-A2

To further explore the binding pattern between peptides and HLA-A2, molecular docking model was established with Galaxypepdock, and pair-wise comparison of pMHC structure was performed between ORF1ab_1707-1716_ and ORF1ab A1708D, ORF1ab_2225-2234_ and ORF1ab I2230T-1, ORF1ab_2230-2238_ and ORF1ab I2230T-2, respectively. It’s observed that these mutations slightly decreased the interaction similarity of mutant peptides to HLA-A2 (Figure 4A). Interestingly, subtle structural alterations of peptides presented by HLA-A2 were observed before and after mutation. Molecular docking comparison between ORF1ab _1707-1716_ (Figure 4B, blue) and ORF1ab A1708D (Figure 4B, red) showed A1708D mutation caused deflection of the benzene ring of the subsequent asparagine (N) (Figure 4C, angle from 145.4 ° to 101.8°). Modeling of ORF1ab_2225-2234_ (Figure 4D, blue) and ORF1ab I2230T-1 (Figure 4D, red) showed that the I2230T mutation herein might affect the later tryptophan (W), making the two benzene rings more convex (Figure 4E, angle from 66.7°to 92.2°). Modeling of ORF1ab_2230-2238_ (Figure 4F, blue) and ORF1ab I2230T-2 (Figure 4F, red) showed that tryptophan (W) was more convex, and its benzene ring tended to expand (Figure 4G, angle from 0 to 66.6 °). These structural changes might provide the possible molecular basis for the altered antigen presentation and CD8^+^ T cell activation, while further protein crystallographic analysis is needed for confirmation.

**Figure 4:**
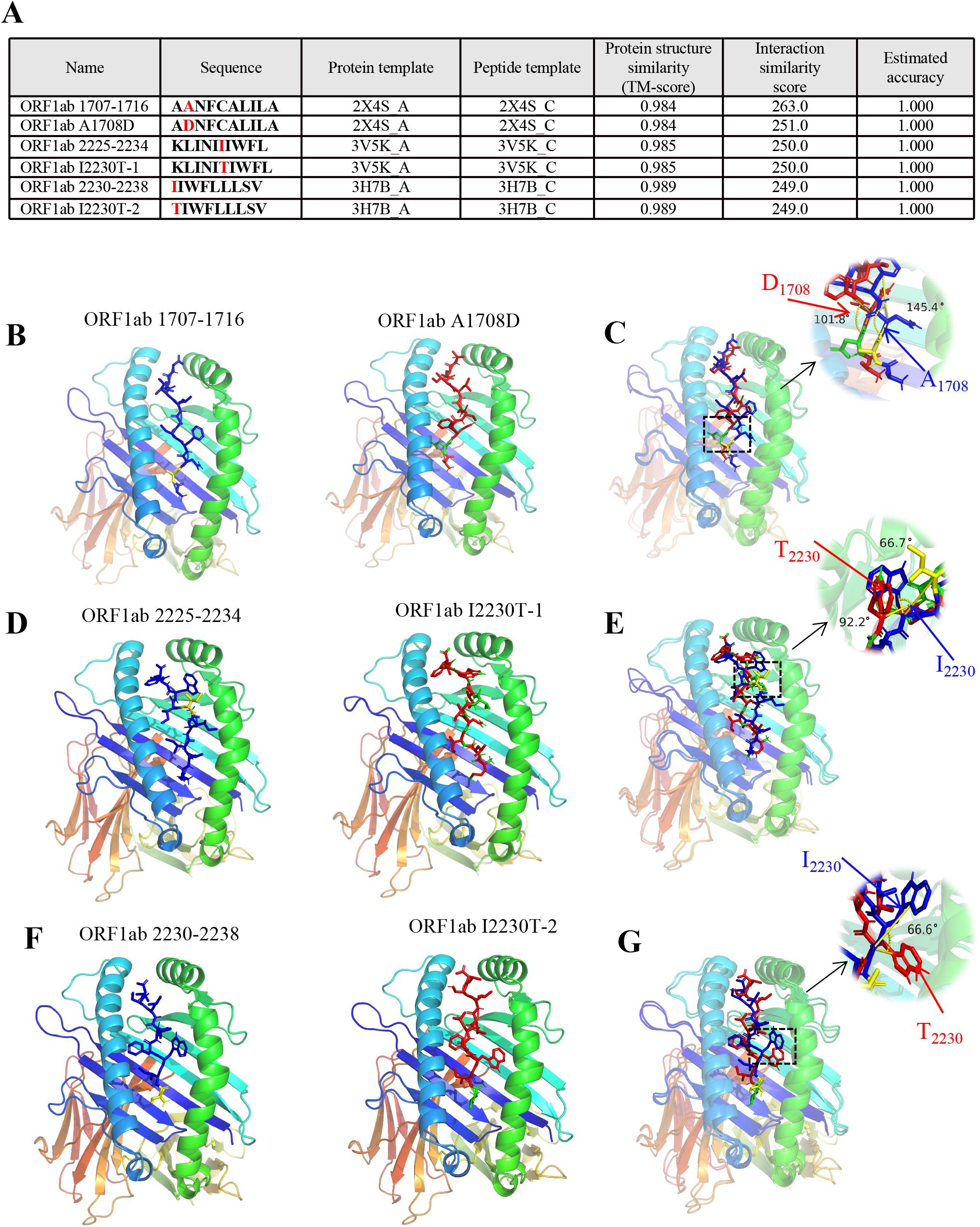
Computational molecular docking simulation of ancestral and mutant antigenic peptides with HLA-A2 molecule. Galaxypepdock was used for molecular docking simulation to demonstrate the structural interaction of HLA-A2 and peptides from ancestral or mutant. **A:** Summary of molecule docking simulation. **B**-**C:** Structures of ORF1ab 1707-1716 (B, blue stick) and ORF1ab A1708D (B, red stick) were compared in B (angle from 145.4°to 101.8°). **D**-**E:** Structures of ORF1ab 2225-2234 (D, blue stick) and ORF1ab I2230T (D, red stick) were compared in D (angle from 66.7°to 92.2°). **F-G:** Structures of ORF1ab 2230-2238 (F, blue stick) and ORF1ab I2230T (F, red stick) were compared in F. (angle from 0 to 66.6°).

## DISCUSSION

The soaring rise of SARS-CoV-2 infection in the last months of 2020 has led to the evolution of several variants with related mutations or characteristics(Lauring and Hodcroft, 2021). One such variant, designated B.1.1.7, was identified in the UK during late 2020 and continued to dominate the circulation in the region. Recent studies have reported longer persistence and higher viral loads in samples from B.1.1.7 infected individuals, indicating its association with the higher infectivity and transmissibility(Calistri et al., 2021; Parker et al., 2021). It’s also reported that B.1.1.7 might even lead to more severe illness(Challen et al., 2021). Our study aimed to fill a key knowledge gap addressing the potential of SARS-CoV-2 variants to evade recognition by human immune responses. Based on the mutation sites in B.1.1.7, we performed computational prediction of HLA-A2-restricted CD8^+^ T cell epitopes, and obtained 19 potential epitopes for ancestral Wuhan strain and 20 for variant B.1.1.7, respectively. To validate the binding of these predicted epitopes, we then checked whether they could be presented on T2A2 cells, where the peptide-MHC complex would be more stabilized if the epitopes bind with HLA-A2 suitably. Our results showed that most of the peptides had reasonable binding with HLA-A2, while the binding capability of most mutant peptides was lower than that of the ancestral. Recently, Tarke et al. reported an identification of 523 CD8^+^ T cell epitopes associated with unique HLA restrictions(Tarke et al., 2021a), and 508 (97.1%) of them were totally conserved within the B.1.1.7 mutant(Tarke et al., 2021b), which might be a reason why they didn’t see significant difference in the T cell reactivity to the ancestral and mutant peptides. By using computational prediction, they reported 73.3% of the mutations were not associated with decrease in binding capacity(Tarke et al., 2021b). The difference of the results may be due to the different verification methods for the binding ability of SARS-CoV-2 T cell epitopes.

Up to date, SARS-CoV-2 mutations of most concern were existed in the viral spike protein, including notable mutations in the receptor binding domain (RBD), N-terminal domain (NTD) and furin cleavage site region. Several of these mutations directly affect ACE2 receptor binding affinity, which may subsequently alter the infectivity, viral load, or transmissibility(Greaney et al., 2021; Zahradnik et al., 2021). Accordingly, it is crucial to address to what extent the mutations from the variants would impact the immunity induced by either SARS-CoV-2 variant infection or vaccination. Currently, most of the studies about immune responses against B.1.1.7 are focusing on alteration of humoral immunity. With neutralization assay to pseudovirus bearing B.1.1.7 spike protein, multiple specific mAbs showed resistance to B.1.1.7 pseudovirus(Collier et al., 2021; Muik et al., 2021). Furthermore, slightly but significantly decreased sensitivity was observed to the sera from SARS-CoV-2 vaccinees and convalescent patients(Muik et al., 2021; Wang et al., 2021). SARS-CoV-2 vaccine clinical trial data demonstrated that specific CD8^+^ response was elicited as well as antibody production(Sahin et al., 2020), and the rapid emergence of the protection at the time when antibodies were still low further supported the important role of cellular immunity(Polack et al., 2020).

So far, the only report assessing the cellular immunity against B.1.1.7 is from Tarke et al, in which they evaluated the CD8^+^ T cell reactivity in convalescent patients by using proteome-wide overlapping peptide megapools, and reported similar responses between ancestral and B1.1.7(Tarke et al., 2021b). In our study, altered CD8^+^ T cell response was observed for particular CD8^+^ epitopes. Our results first did show that mixed epitope-loaded antigen presentation cells could activate T cells from healthy donors. Notably, the proportion of CD8^+^ T cells specific to certain mutant peptides was less than that to ancestral in the same host. In addition, the ancestral epitope specific CD8^+^ T cells could not be recognized by tetramers prepared with mutant epitopes, and vice versa. All these results showed that the T cell mediated immune responses induced by variant B.1.1.7 was decreased. Our previous work has also shown that the L>F mutations in spike protein epitope FVF<u>L</u>VLVPLV resulted in antigen presentation deficiency and reduced specific T cell function, indicating an immune evasion induced by viral evolution(Qiu et al., 2020). The impaired immune responses were further confirmed with the epitope specific CD8^+^ T cells measurement from convalescent COVID-19 patients and SARS-CoV-2 vaccinees. Both results demonstrated that the proportion of T cells recognized by the mutant epitopes of B.1.1.7 was lower than that of the ancestral epitopes in the same individual. In summary, our results indicated that variant B.1.1.7 caused CD8^+^ T cell epitopes mutation, which impaired the CD8^+^ T cell immune response. However, our data showed that the vaccine we used still elicited over 60% of the immune protection against B.1.1.7 based on the cellular immune responses.

Even only focusing on HLA-A2 population, and lacking of information from acute B.1.1.7 infected patients, our data strongly indicated that mutant epitopes in SARS-CoV-2 variant B.1.1.7 caused deficiency in antigen presentation and CD8^+^ T cell immune responses. It’s required to rebuild a new CD8^+^ T cell immune response for variant B.1.1.7.

## Materials and Methods

### Human subjects

The Institutional Review Board of the Affiliated Huaqiao Hospital of Jinan University approved this study. Unexposed donors were healthy individuals enrolled in Guangzhou Blood Center and confirmed with a negative report for SARS-CoV-2 RNA real-time reverse transcriptase polymerase chain reaction (RT-PCR) assay. These donors had no known history of any significant systemic diseases, including, but not limited to, hepatitis B or C, HIV, diabetes, kidney or liver diseases, malignant tumors, or autoimmune diseases. Convalescent donors included subjects who were hospitalized for COVID-19 or confirmed SARS-CoV-2 infection by RT-PCR assay (Supplementary Table 2). SARS-CoV-2 vaccinees were also recruited 1-3 months after vaccination with the inactivated vaccine(Beijing Institute of Biological Products of Sinopharm) (Supplementary Table 3). All subjects provided informed consent at the time of enrollment that their samples could be used for this study. Complete blood samples were collected in acid citrate dextrose tubes and stored at room temperature prior to peripheral blood mononuclear cells (PBMCs) isolation and plasma collection. PBMCs were isolated by density gradient centrifugation using lymphocyte separation medium (GE). After isolation, the cells were cryopreserved in fetal bovine serum (LONSERA) with 10% dimethyl sulfoxide (DMSO) (Sigma-Aldrich) until use.

### HLA-A2 restricted T cell epitope prediction

The spike (S), membrane (M), nucleocapsid (N) and ORF protein sequences of SARS-CoV-2 Wuhan-Hu-1 strain (NC_045512.2) were used for T cell epitope prediction with the “MHC I Binding”tool (http://tools.iedb.org/mhci). The prediction method used was IEDB Recommended 2.22 (NetMHCpan EL), with MHC allele selected as HLA-A*02:01 and HLA-A*02:06, the most frequent class I HLA genotype among Chinese population(González-Galarza et al., 2015; He et al., 2018). All predicted epitopes containing the same amino acid residue corresponding to the mutation from B.1.1.7 were compared. The peptide with the best prediction score was used as the candidate epitope for ancestral Wuhan strain. Meanwhile, peptides with identical amino acid sequences except for the mutated point were used as candidate epitopes for variant B.1.1.7.

### Peptide screening in T2A2 cells

The candidate peptides were synthesized in GenScript Biotechnology Co., Ltd (Nanjing, China) and resuspended in DMSO at a concentration of 10 mM, respectively. T2A2 cells were seeded into 96-well plates, and then incubated with peptides at a final concentration of 20 µM at 37 °C for 4 hours. Set DMSO as blank control, Influenza A M1 peptide (GILGFVFTL) as positive control, and Zika virus peptide (GLQRLGYVL) as negative control. Cells were stained with PE anti-human HLA-A2 antibody (BioLegend) at 4 °C in the dark for 30 min, and acquired in flow cytometer FACS Canto (BD).

### Enzyme-linked immunosorbent assay

10 mM peptide stock solution was diluted to 400 µM in PBS. 20 µL diluted peptide and 20µL 1µg/mL UV-sensitive peptide HLA-A2 monomer (BioLegend) were added into 96-well plates and mixed well by pipetting up and down. The plates were then exposed to UV light (365 nm) for 30 min on ice, and incubated for 30 min at 37 °C in the dark. Finally, 40 µL of peptide-exchanged monomer was used for test. The pMHC level was determined by using a pMHC ELISA Kit (Mlbio). Within 15 min after adding stop solution, the absorbance values of sample were read at 450 nm. Set UV-irradiated monomers as blank control, Influenza A M1 peptide (GILGFVFTL) as positive control, and Zika virus peptide (GLQRLGYVL) as negative control.

### Generation of antigen specific HLA-A2 tetramer

30 µL peptide-exchanged monomer formed in the above steps was mixed with 3.3 µL PE streptavidin (BioLegend) on a new plate and incubated on ice in the dark for 30 min. 2.4 µL blocking solution (1.6 µL 50mM biotin plus 198.4 µL PBS) was added to stop the reaction and incubated at 4-8 °C overnight.

### Cell-surface CD8, CCR7, CD45RA and tetramer staining

PBMCs were isolated from peripheral venous blood of healthy donors, convalescent COVID-19 patients and SARS-CoV-2 vaccinees. The HLA-A2^+^ donors were identified by using flow cytometry. Briefly, 10^6^ PBMCs were stained with FITC anti-human HLA-A2 antibody (BioLegend) at 4 °C in the dark for 30 min, and acquired by using flow cytometer. HLA-A2 positive PBMC samples were further stained with PE labeled tetramer (home-made), PerCP labeled human CD8^+^ antibody (BioLegend), APC labeled human CCR7 antibody (BioLegend), FITC labeled human CD45RA antibody (BioLegend) and acquired with flow cytometer FACS Canto (BD).

### Activation and cytotoxicity analysis of CD8^+^ T cells

HLA-A2 expressing T2A2 cells were loaded with peptides for subsequent T cell activation. Briefly, T2A2 cells were treated with 20 µg/mL mitomycin C (Sinochem) for 30 min to stop cell proliferation, and loaded with given epitope peptides. 0.5×10^6^ CD8^+^ T cells isolated from health donors were co-cultured with 0.5 × 10^6^ peptide-loaded T2A2 cells stained with 5 µmol/L CFSE (TargetMol), and stimulated with 1 µg/mL anti-human CD28 antibodies (BioLegend) and 50 IU/mL IL-2 (SL PHARM, Recombinant Human Interleukin-2(^125^Ala) Injection). 50 IU/mL IL-2 and 20 µM mixed peptides were then supplemented every two days. The T cell activation marker CD69 (BioLegend), tetramer specific CD8^+^ T cells and apoptosis marker Annexin V-APC (BioLegend) on T2A2 cells were evaluated after 16 hours and 7 days, respectively. On day 7, cells were re-stimulated with peptides for 6 hours in the presence of Leuko Act Cktl with GolgiPlug (BD) plus 50 IU/mL IL-2, and the production of IFN-γ and Granzyme B was checked with PerCP anti-human IFN-γ (BioLegend) and FITC anti-human Granzyme B (BioLegend) staining.

### Molecular Docking simulation of peptide-HLA-A2 complex

To evaluate the binding pattern and affinity of peptides with HLA-A2, molecular docking simulation was carried out with Galaxypepdock. The available structure of HLA: 0201 (PDB ID: 3mrb) was downloaded from the RSCB PDB server (https://www.rcsb.org/) for modeling. Galaxypepdock is a template-based docking program for peptides and proteins, which can generate 10 models to evaluate the results of the docking(Hasup et al., 2015; Mani et al., 2020). The top model with the highest interaction similarity score was selected and visualized by using Discovery Studio 4.5. PyMol 1.1 software was used to calculate the angle deflection of benzene ring in the polypeptide, and used the central atoms of three amino acids to calculate the angle.

### Statistical Analysis

The data were analyzed by one-way ANOVA and paired-samples t-tests for statistical significance by using Graphpad prism 8 and SPSS 22.0 software. *P* value less than 0.05 was considered to be statistically significant.

## Acknowledgements

This work was supported by grants from the National Key Research and Development Program of China (2018YFC2002003), the Natural Science Foundation of China (U1801285, 81971301), Guangzhou Planned Project of Science and Technology (201904010111, 202002020039), Zhuhai Planned Project of Science and Technology (ZH22036302200067PWC) and the Initial Supporting Foundation of Jinan University.

## Author contributions

G.C., P.W. and L.X. designed the project. C.X. performed the experiments. L.M. performed the molecular docking simulation. Z.W., G.Z. and Z.Y. analyzed the clinical information and performed sample collection. L.G., and J.S. assisted with experiments; X.C., J.Y., Y.H., J.X., W.J. H.N. and F.G. assisted with clinical information and sample collection. C.Q., O.J.L., P.W. and G.C. analyzed the data. O.J.L. assisted with data analysis. C.X., P.W. and G.C. wrote the manuscript.

## Supplementary information

### Supplementary Figures

Supplementary Figure 1: Identification of immunogenicity of HLA-A2 restricted T cell epitope of SARS-CoV-2 variant B.1.1.7

### Supplementary Tables

Supplementary Table 1: Raw data of predicted CD8^+^ T cell epitopes for SARS-CoV-2 variant B.1.1.7

Supplementary Table 2: Clinical characteristics of convalescent COVID-19 patients

Supplementary Table 3: Clinical characteristics of SARS-CoV-2 vaccinees

### Supplementary data set

Raw data of protein docking, PDB files.

