## Supplementary Figure 1 for "SARS-CoV-2 variant B.1.1.7 caused HLA-A2^+^ CD8^+^ T cell epitope mutations for impaired cellular immune response"

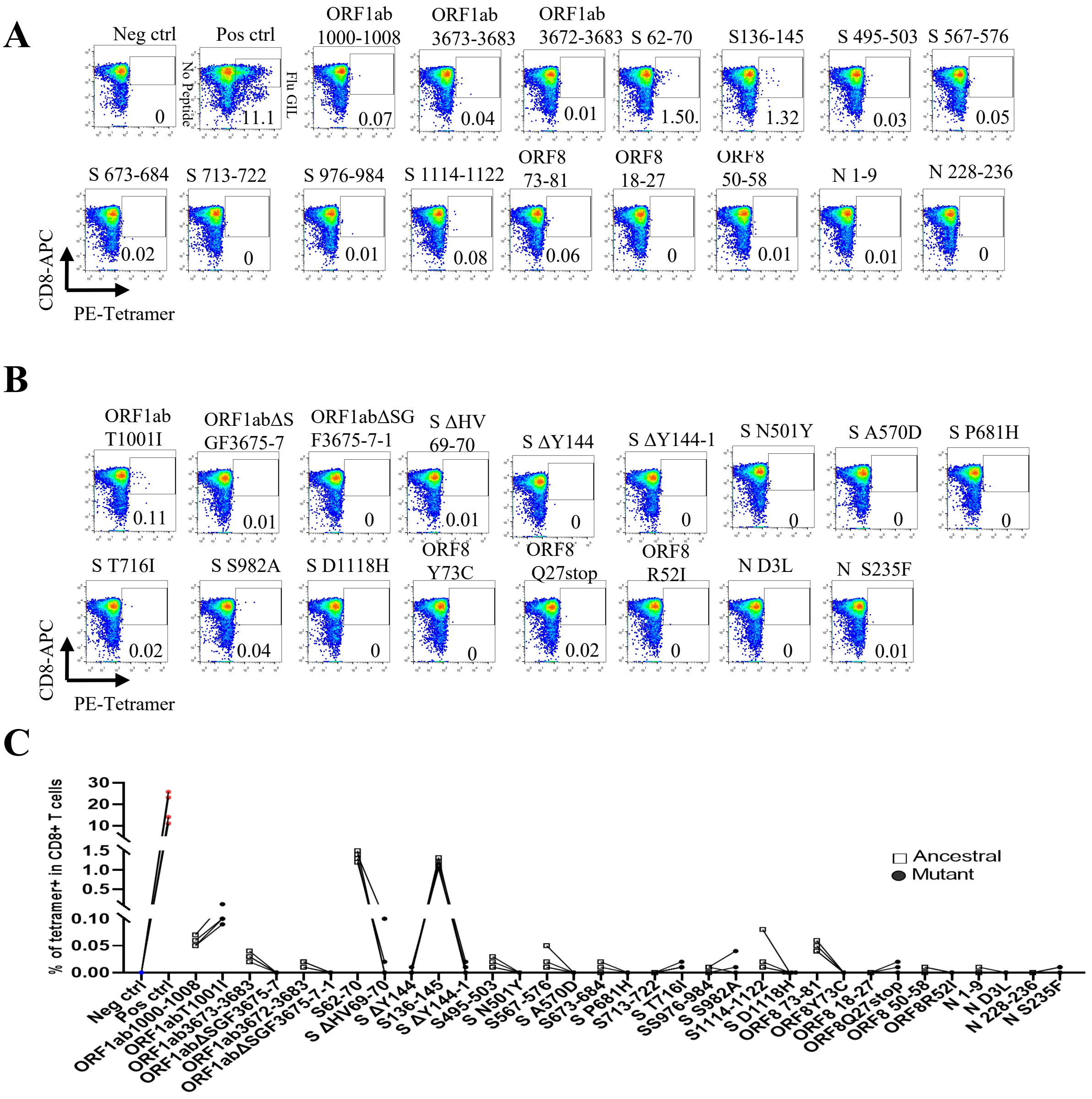


### Figure Legends

**Figure S1 Identification of immunogenicity of HLA-A2 restricted T cell epitope of SARS-CoV-2 variant B.1.1.7**

Mitomycin pretreated T2A2 cells were loaded with mixed peptides from ancestral or mutant, and incubated with CD8+ T cell from health donors at 1:1 ratio, respectively. Epitope specific CD8+ T cells were generated after 7 day stimulation.

**A-C:** In the same healthy volunteers, the ancestral(A) and mutated (B) epitopes near the mutation site stimulated healthy CD8+ T cells to generate antigen-specific T cells for 7 days. C was the representative plot of A and B (Symbols represent individual person, n = 4 per group).Ancestral: Wuhan strain epitope; Mutant: varian B.1.1.7 epitope. Neg ctrl: T2A2 without peptide loading; Pos ctrl: T2A2 loaded with influenza A M1 peptide GILGFVFTL.
