## Supplementary Table 1 for "SARS-CoV-2 variant B.1.1.7 caused HLA-A2^+^ CD8^+^ T cell epitope mutations for impaired cellular immune response"

| **Table S1** Raw data of predicted CD8+ T cell epitopes for SARS-CoV-2 variant B.1.1.7 | | | | | | | | | | | | | | | | |  |
| --- | --- | --- | --- | --- | --- | --- | --- | --- | --- | --- | --- | --- | --- | --- | --- | --- | --- |
| Protein | Allele | Seq_num | Start | End | Length | Sequence | Method | Percentile | Rank | ann_ic50 | ann_rank | smm_ic50 | smm_rank | comblib_sidney2008_score | comblib_sidney2008_rank | netmhcpan_ic50 | netmhcpan_rank |
| **ORF1ab** | HLA-A*02:06 | 1 | 1000 | 1008 | 9 | TTIQTIVEV | Consensus | (ann/smm) | 1.39 | 22.3 | 0.27 | 77.37 | 2.5 | - | - | - | - |
|  | HLA-A*02:06 | 1 | 1707 | 1716 | 10 | AANFCALILA | Consensus | (ann/smm) | 9.55 | 359.03 | 2.1 | 1210.35 | 17 | - | - | - | - |
|  | HLA-A*02:01 | 1 | 2225 | 2234 | 10 | KLINIIIWFL | Consensus | (ann/smm) | 0.13 | 7.29 | 0.05 | 6.45 | 0.2 | - | - | - | - |
|  | HLA-A*02:01 | 1 | 2230 | 2238 | 9 | IIWFLLLSV | Consensus | (ann/comblib_sidney2008/smm) | 0.4 | 19.01 | 0.21 | 18.84 | 0.4 | 2.15E-04 | 12 | - | - |
|  | HLA-A*02:01 | 1 | 3673 | 3683 | 11 | SLSGFKLKDCV | Consensus | (ann/smm) | 4.55 | 2410.71 | 7.7 | 166.31 | 1.4 | - | - | - | - |
|  | HLA-A*02:01 | 1 | 3672 | 3683 | 12 | TSLSGFKLKDCV | Consensus | (ann/smm) | 9.4 | 3380.3 | 9.4 | - | - | - | - | - | - |
| **Spike** | HLA-A*02:06 | 1 | 62 | 70 | 9 | VTWFHAIHV | Consensus | (ann/smm) | 1.02 | 74.83 | 0.75 | 37.89 | 1.3 | - | - | - | - |
|  | HLA-A*02:01 | 1 | 136 | 145 | 10 | CNDPFLGVYY | Consensus | (ann/smm) | 43 | 25302.05 | 44 | 21952.3 | 42 | - | - | - | - |
|  | HLA-A*02:06 | 1 | 495 | 503 | 9 | YGFQPTNGV | Consensus | (ann/smm) | 2.6 | 736.15 | 3.2 | 59.23 | 2 | - | - | - | - |
|  | HLA-A*02:01 | 1 | 567 | 576 | 10 | RDIADTTDAV | Consensus | (ann/smm) | 0.03 | 845 | 8 | - | 2.5 | - | - | - | - |
|  | HLA-A*02:06 | 1 | 673 | 684 | 12 | SYQTQTNSPRRA | Consensus | (ann/smm) | 14306.68 | 28 | - | - | - | - | - | - |  |
|  | HLA-A*02:06 | 1 | 713 | 722 | 10 | AIPTNFTISV | Consensus | (ann/smm) | 1.4 | 118.96 | 1.2 | 89.93 | 1.6 | - | - | - | - |
|  | HLA-A*02:01 | 1 | 976 | 984 | 9 | VLNDILSRL | Consensus | (ann/comblib_sidney2008/smm) | 1.1 | 33.57 | 0.38 | 58.36 | 1.1 | 9.73E-05 | 5.5 | - | - |
|  | HLA-A*02:06 | 1 | 1114 | 1122 | 9 | IITTDNTFV | Consensus | (ann/smm) | 2.35 | 131.91 | 1.3 | 104.6 | 3.4 | - | - | - | - |
|  |  |  |  |  |  |  |  |  |  |  | - | - | - | - | - | - | - |
| **ORF8** | HLA-A*02:06 | 1 | 73 | 81 | 9 | YIDIGNYTV | Consensus | Consensus | 0.957895 | 0.01 | - | - | - | - | - | - | - |
|  | HLA-A*02:06 | 1 | 18 | 27 | 10 | QECSLQSCTQ | Consensus | Consensus | 1.50E-05 | 84 | - | - | - | - | - | - | - |
|  | HLA-A*02:06 | 1 | 50 | 58 | 9 | GARKSAPLI | Consensus | Consensus | 0.004488 | 15 | - | - | - | - | - | - | - |
| **N** | HLA-A*02:06 | 1 | 1 | 9 | 9 | MSDNGPQNQ | Consensus | Consensus | 0.000733 | 22 | - | - | - | - | - | - | - |
|  | HLA-A*02:06 | 1 | 228 | 236 | 9 | NQLESKMSG | Consensus | (ann/smm) | 14 | 7010.51 | 16 | 462.97 | 12 | - | - | - | - |
