## Supplementary Table 2 for "SARS-CoV-2 variant B.1.1.7 caused HLA-A2^+^ CD8^+^ T cell epitope mutations for impaired cellular immune response"

**Table S2** Clinical characteristics of convalescent COVID-19 patients

| Donor code | SARS CoV-2  PCR | Data of sample collection | Sex | Age | HLA-A2 restriction | Symptoms | % of tetramer+ cells | | | | | | | |
| --- | --- | --- | --- | --- | --- | --- | --- | --- | --- | --- | --- | --- | --- | --- |
|  |  |  |  |  |  |  | ORF1ab  1707-1716 | ORF1ab  A1708D | ORF1ab  2225-2234 | ORF1ab  I2230T-1 | | ORF1ab  2230-2238 | | ORF1ab  I2230T-2 |
| Case 1 | Positive | 2020/4/12 | Male | 54 | Yes | - | 7.15 | 4.67 | 37.6 | 11.2 | 8.92 | | 5.89 | |
| Case 2 | Positive | 2020/5/20 | Female | 31 | No | - | - | - | - | - | - | | - | |
| Case 3 | Positive | 2020/5/20 | Female | 38 | No | - | - | - | - | - | - | | - | |
| Case 4 | Positive | 2020/5/20 | Female | 53 | No | - | - | - | - | - | - | | - | |
| Case 5 | Positive | 2020/6/8 | Male | 50 | No | - | - | - | - | - | - | | - | |
| Case 6 | Positive | 2020/6/8 | Female | 23 | No | - | - | - | - | - | - | | - | |
| Case 7 | Positive | 2020/6/17 | Male | 35 | No | - | - | - | - | - | - | | - | |
| Case 8 | Positive | 2020/6/17 | Male | 30 | No | - | - | - | - | - | - | | - | |
| Case 9 | Positive | 2020/6/17 | Male | 31 | No | - | - | - | - | - | - | | - | |
| Case 10 | Positive | 2020/6/17 | Male | 29 | No | - | - | - | - | - | - | | - | |
| Case 11 | Positive | 2020/6/17 | Male | 35 | No | - | - | - | - | - | - | | - | |
| Case 12 | Positive | 2020/09/19 | Male | 64 | No | Mild | - | - | - | - | - | | - | |
| Case 13 | Positive | 2020/09/19 | Female | 54 | No | Mild | - | - | - | - | - | | - | |
| Case 14 | Positive | 2020/09/19 | Female | 63 | No | Mild | - | - | - | - | - | | - | |
| Case 15 | Positive | 2020/10/18 | Female | 66 | No | Mild | - | - | - | - | - | | - | |
| Case 16 | Positive | 2020/10/18 | Female | 57 | Yes | Mild | 4.28 | 3.04 | 9.67 | 7.97 | 7.17 | | 6.48 | |
| Case17 | Positive | 2020/10/18 | Female | 54 | Yes | Mild | 3.49 | 3.22 | 11.2 | 10.3 | 7.7 | | 5.32 | |
| Case 18 | Positive | 2020/10/23 | Male | 61 | No | Mild | - | - | - | - | - | | - | |
| Case 19 | Positive | 2020/10/23 | Female | 57 | Yes | Mild | 3.69 | 2.94 | 10.2 | 7.06 | 8.37 | | 6.2 | |
| Case 20 | Positive | 2020/11/09 | Female | 74 | No | Mild | - | - | - | - | - | | - | |
| Case 21 | Positive | 2020/11/09 | Female | 64 | No | Mild | - | - | - | - | - | | - | |
| Case 22 | Positive | 2020/11/09 | Female | 49 | No | Mild | - | - | - | - | - | | - | |
| Case 23 | Positive | 2020/12/06 | Female | 77 | No | Severe | - | - | - | - | - | | - | |
| Case 24 | Positive | 2020/12/06 | Male | 60 | No | Severe | - | - | - | - | - | | - | |
| Case 25 | Positive | 2020/12/17 | Female | 65 | No | Severe | - | - | - | - | - | | - | |
