## Supplementary Table 3 for "SARS-CoV-2 variant B.1.1.7 caused HLA-A2^+^ CD8^+^ T cell epitope mutations for impaired cellular immune response"

**Table S3** Clinical characteristics of SARS-CoV-2 vaccinees

| Donor code | HLA-A2  restriction | Sex | Age | BMI  (kg/m^2^) | First vaccination time | Time for booster immunization | Blood collection time | Symptoms |
| --- | --- | --- | --- | --- | --- | --- | --- | --- |
| # 1 | Yes | Female | 22 | 19.83 | 2020/9/3 | 2020/9/25 | 2021/1/23 | NO |
| # 2 | Yes | Female | 36 | 18.49 | 2020/9/3 | 2020/9/25 | 2021/1/23 | NO |
| # 3 | Yes | Female | 33 | 20.57 | 2020/9/3 | 2020/9/25 | 2021/1/23 | NO |
| # 4 | Yes | Female | 51 | 25.63 | 2020/9/3 | 2020/9/25 | 2021/1/23 | Local redness and swelling |
| # 5 | Yes | Male | 55 | 25.35 | 2020/9/3 | 2020/9/25 | 2021/1/23 | NO |
| # 6 | Yes | Male | 58 | 22.49 | 2020/9/3 | 2020/9/25 | 2021/1/23 | NO |
| # 7 | Yes | Female | 28 | 20.76 | 2020/9/3 | 2020/9/25 | 2021/1/23 | NO |
| # 8 | Yes | Female | 53 | 20.43 | 2020/9/3 | 2020/9/25 | 2021/1/23 | Dizziness for  4 hours |
| # 9 | Yes | Male | 21 | 22.49 | 2020/12/23 | 2021/1/18 | 2021/1/23 | NO |
| # 10 | Yes | Male | 22 | 23.39 | 2020/12/23 | 2021/1/18 | 2021/1/23 | NO |
| # 11 | Yes | Male | 38 | 24.42 | 2020/12/24 | 2021/1/18 | 2021/1/23 | NO |
| # 12 | Yes | Male | 22 | 22.94 | 2020/12/23 | 2021/1/20 | 2021/1/23 | NO |
| # 13 | Yes | Female | 42 | 23.07 | 2020/9/25 | 2020/9/25 | 2021/1/23 | NO |
| # 14 | Yes | Male | 39 | 27.78 | 2020/9/25 | 2020/9/25 | 2021/1/23 | NO |
| # 15 | Yes | Female | 53 | 25.34 | 2020/9/25 | 2020/9/25 | 2021/1/23 | NO |
| # 16 | Yes | Female | 21 | 19..36 | 2020/12/23 | 2021/1/18 | 2021/1/23 | NO |
| # 17 | Yes | Male | 43 | 23.14 | 2020/12/23 | 2021/1/18 | 2021/1/23 | NO |
